# A Dynamical Density Functional Theory Framework for Non-Equilibrium Phase Dynamics in Biomolecular Condensates

**DOI:** 10.1101/2025.10.02.680159

**Authors:** Alexander R.J. Silalahi, Morgan G. Murray, Gül H. Zerze

**Affiliations:** Chemical and Biomolecular Engineering Department, University of Houston

**Author notes:** (Electronic mail) (Gül H. Zerze).

## Abstract

Biomolecular condensates, membrane-less organelles that arise from liquid-liquid phase separation (LLPS) of proteins and nucleic acids, play vital roles in cellular organization and regulation. Computational modeling is crucial for uncovering the molecular mechanisms behind LLPS; however, the fluctuations across a wide range of spatial scales and the inherently non-equilibrium nature of these systems make capturing their long-timescale dynamics particularly challenging. Here, we present a continuum dynamical density functional theory (DDFT) framework that captures the non-equilibrium dynamics of LLPS by integrating a physics-based statistical mechanical theory with key experimentally-derived parameters. Our model couples DDFT with a continuum free energy functional, incorporating two-body correlations between monomers and surface tension effects to determine binodal densities under phase coexistence. By solving the DDFT equations, we describe the time evolution of phase-separated domains, capturing key long-timescale processes such as droplet maturation, coalescence, and interface relaxation, phenomena that are difficult to probe using atomistic or mesoscale coarse-grained simulations. This implementation integrates experimental phase equilibrium data with molecular-scale descriptors, such as amino acid properties, to construct a quantitative link between molecular interactions and macroscopic phase behavior. However, the approach is generalizable, providing a foundation for self-contained molecular-to-continuum modeling bridge platforms.

## I. INTRODUCTION

Biomolecular condensates formed via liquid–liquid phase separation (LLPS) underlie diverse cellular processes, including transcription, stress response, and compartmentalization of biochemical reactions^1,2^. Understanding the dynamics of their formation, dissolution, and morphological evolution is key to deciphering their biological roles and disease connections^2–6^. However, the timescales over which these behaviors unfold span orders of magnitude—from microseconds of molecular reorganization to minutes and hours of droplet coarsening and stabilization—posing challenges for both experiments and simulations.

Experimental techniques such as fluorescence microscopy, optical trapping, and microrheology have advanced our understanding of the physical mechanisms underlying the condensate formation^1,7–10^. Although these experimental methods are highly effective for characterizing condensate morphology, material properties, and dynamics across physiologically relevant timescales, directly resolving molecular mechanisms, such as the contribution of specific sequence features, transient interactions, or local compositional heterogeneity, often requires indirect inference with these techniques. Computational models provide a valuable complement by offering molecular-scale perspectives on the thermodynamics and kinetics of phase separation, helping to interpret and contextualize experimental findings^3,11,12^.

Molecular simulations, including atomistic and coarse-grained molecular dynamics (MD), have been extensively used to study the microscopic driving forces of phase separation^11–17^. These approaches provide detailed information on protein-protein and protein-RNA interactions, conformational dynamics, and sequence effects. However, they are limited in accessible length and timescales^18^, making it challenging to capture long-term evolution and coarsening of condensates.

In parallel to molecular simulations and experiments, continuum modeling approaches such as Flory-Huggins (FH) theory^19,20^, Cahn-Hilliard phase field models^21^, and reaction-coupled phase separation models^22–24^ have been employed to study biomolecular LLPS^11,12,25,26^. These models offer a thermodynamic perspective on the continuum scale, allowing for the description of phase separation in terms of macroscopic parameters such as binodal curves, interfacial tension, and coarsening dynamics. In particular, FH theory is often used to extract effective interaction parameters (*χ*) from experiments or molecular simulations. Phenomenology-based phase-field models and Cahn-Hilliard models describe the time evolution of phase-separated domains on larger length and timescales. These models are appealing due to their scalability and simplicity, but they rely on empirically fitted free energy functions, which can limit predictive power and hinder connections to underlying molecular interactions.

To address this limitation, here we adopted a dynamical density functional theory (DDFT) framework that enables a physics-grounded, simulation-compatible approach to modeling the long-timescale behavior of condensates. Derived from statistical mechanical principles, DDFT provides a systematic route to encode interaction potentials and concentration-dependent correlations into continuum dynamics. Its modular structure allows the incorporation of interaction-specific terms, such as mean-field attraction, electrostatics, and interfacial effects—offering a path toward bridging molecular simulations with macroscopic observables.

In our current implementation, we incorporated Flory–Huggins (FH)-like interaction parameters, random phase approximation (RPA)-based electrostatic interactions^27–29^, and surface tension terms into the free energy functional. The FH parameters were obtained by fitting experimental phase equilibrium data, introducing phenomenological elements into the model. However, the same parameters can also be obtained from phase diagrams generated via molecular simulations, enabling future simulation-based multiscaling platforms. Thus, the framework retains its physics-based foundation while allowing flexibility in input sources—providing a scalable and extensible platform for modeling the equilibrium and dynamic features of biomolecular phase separation.

## II. METHODS

Continuum modeling approaches for phase separation in intrinsically disordered protein (IDP) systems often build on polymer physics frameworks, where proteins are treated as flexible chains of residues interacting via sequence-specific forces in solution. In that representation, each monomer corresponds to an amino acid, and the system is embedded in an aqueous environment with small counterions and possibly added salt^30,31^. The equilibrium behavior of such systems, such as critical temperature, binodal curves, and concentration-dependent miscibility, is commonly captured using mean-field theories or field-theoretic polymer models, where the total free energy is decomposed into entropic and interaction terms.

Previous field-theoretic studies^32–35^ have modeled equilibrium LLPS behavior by incorporating electrostatic interactions and charge patterning at the residue level. For instance, Lin and co-workers ^30^ used a random phase approximation (RPA)-based framework to correlate single-chain compactness with phase separation propensity in charged IDPs. In RPA, the electrostatic component of the free energy captures effective two-body interactions between charged residues and can be extended to account for ionic strength. Within this theoretical construct, the contributions to the overall energy can be decoupled into entropic and electrostatic free energy components.

An additional free energy term to account for *π*-cation and *π*-*π* bond interactions between residues is also included to improve agreement with experimental data^30^. These models successfully predict how sequence charge distribution and salt concentration influence the critical temperature and binodal compositions. However, these models focus on studying the equilibrium predictions and do not describe the spatial and temporal evolution of phase-separated compartments. The dynamics associated with droplet formation or coalescence present a distinct challenge. The goal of this work is to incorporate equilibrium analytical models into a dynamical density functional theory (DDFT) framework capable of simulating the spatiotemporal evolution of condensates.

### A. Dynamical Density Functional Theory (DDFT)

In continuum descriptions of soft matter systems, the local number density is treated as a coarse-grained field that characterizes the system’s configuration. In dynamical density functional theory (DDFT), the system’s time evolution is governed by a continuity equation in which the flux is driven by gradients of the chemical potential derived from a free energy functional. Although this functional is typically borrowed from equilibrium statistical mechanics, DDFT applies it under the adiabatic approximation^36^, i.e., assuming that at each instant the non-equilibrium correlations of the system can be approximated by those of an equilibrium system with the same instantaneous density distribution. This allows DDFT to model nonequilibrium dynamics while retaining molecular-scale physical detail from equilibrium formulations.

The time evolution of the density is driven by the chemical potential derived from the free energy of the corresponding quasi-equilibrium state. This chemical potential serves as the driving force for the density dynamics, guiding the system through a sequence of quasi-equilibrium states as it evolves over time.

The DDFT equation for the density of species *i* (*ρ*_*i*_) is given by:

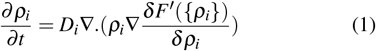

where 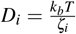 is the diffusion constant of the *i*^*th*^ species in the system. In our current formulation, we only solve this equation for a single protein species. The left term represents the change of density as a function of time, and the right side is the driving force for dynamical change that depends on the functional form of free energy.

We express the free energy as a functional of the volume fraction. Specifically, we define the volume fraction of species *i* as φ_*i*_ = *ρ*_*i*_*v*_*i*_, where *v*_*i*_ = *r*_*i*_*a*^3^ is the volume of a monomer of species *i*, with *a*^3^ being the volume of a lattice site and *r*_*i*_ a dimensionless ratio accounting for monomer size relative to the lattice unit. For simplicity, and to maintain consistency with the lattice-based discretization used in our model, we assume *r*_*i*_ = 1, yielding φ_*i*_ = *ρ*_*i*_*a*^3^. This formulation enables us to express the free energy in terms of volume fractions as follows:

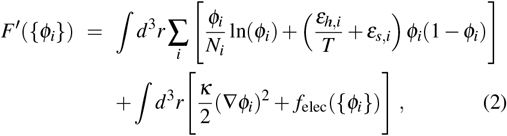

where *N*_*i*_ is the number of monomers in species *i, ε*_*h,i*_ and *ε*_*s,I*_ are fitted Flory Huggins parameters, *κ* is the influence parameter (surface tension related parameter, also see the *Surface Tension* subsection), and *f*_*elec*_ is the electrostatic interactions that take two-body correlations into account. The first integral accounts for the entropic and enthalpic contributions to mixing (including Flory-Huggins-like terms) and the second integral includes interfacial energy (introduced as the density gradient theory^37^) and long-range interactions such as electrostatics (as derived from the RPA model developed by Chan and coworkers^27,30^).

The chemical potential of species *i* can then is then derived from equation 2 as:

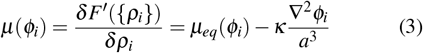

where *µ*_*eq*_ is the contribution to the chemical potential from the free energy terms other than surface tension. The derivations related to the surface tension term are shown below. The derivations of the component of *µ*_*eq*_ are given in Appendix A.

A key limitation of DDFT is that it does not capture the hydrodynamic forces that arise from fluid motion and momentum transport, which can play an important role in the dynamics and coarsening behavior of phase-separating systems. However, we argue that because the system is small enough (micron-scale) in these liquid droplets, surface tension dominates the hydrodynamics^38–40^.

### B. Surface Tension

We first estimated the surface tension contribution arising from the density gap observed in the two-phase coexistence at equilibrium. This surface tension term accounts for the interfacial forces that govern the behavior of the system at the boundary between the dense and dilute phases.

Subsequently, we incorporated this surface tension contribution into the free energy functional that explicitly captures the spatial variation of density. By accounting for the non-uniform distribution of density across different regions of the system, this formulation enables the modeling of the dynamic evolution from the initial density distribution towards the equilibrium density profile. The incorporation of surface tension contributions in free energy formulation is a well-studied and established approach in the literature, commonly referred to as density gradient models^37^.

In the case of a planar interface between dense and dilute phases, surface tension *γ* can be derived by minimizing the total free energy density with respect to density, resulting in (see Appendix B for the derivation)

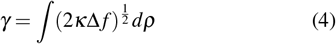

where *κ* is called the influence parameter, *f* is the free energy density, *ρ* is the density of the liquid, and Δ *f* is the difference in the free energy density between a homogeneous mixture of liquid and droplets.

Equation 4 was derived with the assumption that the contribution of the surface tension energy comes from the gradient of density in space and is given by:

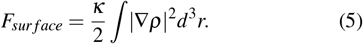

The magnitude of the surface tension can be empirically estimated using the well-established Parachor model^41,42^:

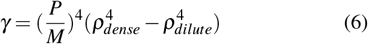

where *γ* is the surface tension at the planar interface that separates the dilute and dense phases of densities *ρ*_*dilute*_ and *ρ*_*dense*_. In this empirical formulation, *P* is the parachor number derived from the polymer constituents and *M* is the polymer’s molar mass. Using this experimentally determined *γ*, we can write eqn 5 with *κ* as follows:

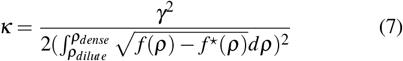

Where

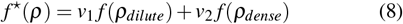

describes the free energy density of the system with two phases with weight factors *v*_1_ and *v*_2_, which are given by:

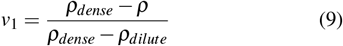

And

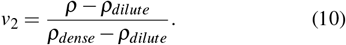

Here, *f* (*ρ*) describes the free energy density of a system with density *ρ*. In this paper, we assume that *κ* is constant for all values of *ρ*. We will explore models in which *κ* takes a *ρ*-dependent form in future work. Further details regarding the influence parameter derivation are provided in Appendix B.

### C. Numerical Solution

Equations 1,2, and 3 serve as the equations governing the dynamics of liquid droplets. We solved these equations using the Finite Volume Method, using `FiPy`^43^ and `Trilinos`^44^ packages. We discretized Eq. 3 using a uniform 2D Cartesian grid with grid spacing Δ*x* = Δ*y* = 1.0Å. Due to the highly nonlinear nature of the governing equations, time integration was performed using an implicit Euler scheme. This implicit time-stepping method is more robust and stable compared to explicit schemes, particularly for nonlinear systems and stiff differential equations. The time step Δ*t* is chosen adaptively within the bound 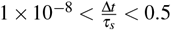.

The cellular or test tube environment that facilitates the formation of liquid droplets is typically much larger than the size of individual proteins. Simulating the droplet and its environment demands an immense computational cost, making it intractable. To overcome this challenge, we divided the computational domain into two regions: i) a domain representing the interactions between liquid droplets and ii) a reservoir domain representing the surrounding environment in which the liquid density is maintained at a constant value. This approach allows for more efficient simulations while capturing the system’s essential dynamics.

We implemented our approach in 2D for initial exploration, due to its computational efficiency, which allowed for the examination of the parameter space while capturing the essential physics of droplet growth and coalescence. We will have the 3D implementation in our future work.

A Robin boundary condition was implemented at the interface between these two domains. This boundary condition allows for the exchange of material across the boundary, with the flux being proportional to the difference in density between the two domains. We consider this formulation to be more physical for our case than insulating (Neumann) or fixed-value (Dirichlet) boundary conditions. Mathematically, the Robin boundary condition can be expressed as:

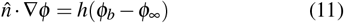

which is implemented for eqn 1 at the boundary, where φ_*b*_ and φ_∞_ are densities at the boundary and of the surrounding environment, respectively, and the magnitude of *h* represents the rate of exchange of liquid phase at the boundary. Robin boundary condition has previously been used in the context of the Cahn-Hillard formalism^45–47^.

## III. RESULTS AND DISCUSSIONS

Here, we used the DDFT framework to investigate the coalescence and relaxation dynamics of condensate droplets composed of Ddx4^N1^, an intrinsically disordered protein (IDP) with a net charge of −4*e* and sequence length *N* = 236 (the sequence is given in Supplementary Material). These simulations explore how interfacial tension, droplet size, and environmental exchange govern the timescale and morphology of droplet coarsening and relaxation.

All DDFT simulations used the free energy functional described in Eq. 2, with Flory-Huggins parameters previously fit to match experimental phase diagrams of Ddx4^N127,30^, namely *ε*_*h*_ = 0.15 and *ε*_*s*_ = −0.3. We also took the monomer and counterions to have the same volume as water molecules *r*_*m*_ = *r*_*c*_ = 1. We adopted a two-dimensional computational domain discretized into a uniform 2D Cartesian mesh. The dimensions *x* and *y* have units of Å. The governing equations are numerically solved using a finite-volume method. DDFT simulations were performed at dimensionless temperature 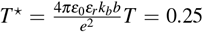. The given FH parameters at the given reduced temperature yield to liquid-liquid phase coexistence with volume fraction (φ_*m*_) of dilute and dense phases of 3.8·10^−4^ and 0.142, respectively, at equilibrium. In the following results, the droplets are initially set to have φ_*m*_ = 0.127 of dense phases.

To interpret the simulation timescales, we estimate the diffusivity of Ddx4^N1^ using the Zimm model, appropriate for flexible polymers in dilute solution^48^:

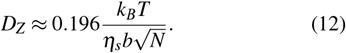

Here, *b* = 3.8 Å is the monomer spacing, *η*_*s*_ = 10^−3^ Pa s is solvent viscosity, and *N* = 236 is the number of amino acids. This gives a characteristic time unit 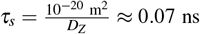, which sets the timescale of DDFT evolution.

We first simulated the coalescence of two adjacent droplets initialized with a uniform dense-phase volume fraction φ = 0.127, which is 90% of the equilibrium dense phase reported in Lin et al.^30^. Figure 1 and 2 show two representative cases with different surface tension parameters (*κ* = 20 and 200).

**FIG. 1.**
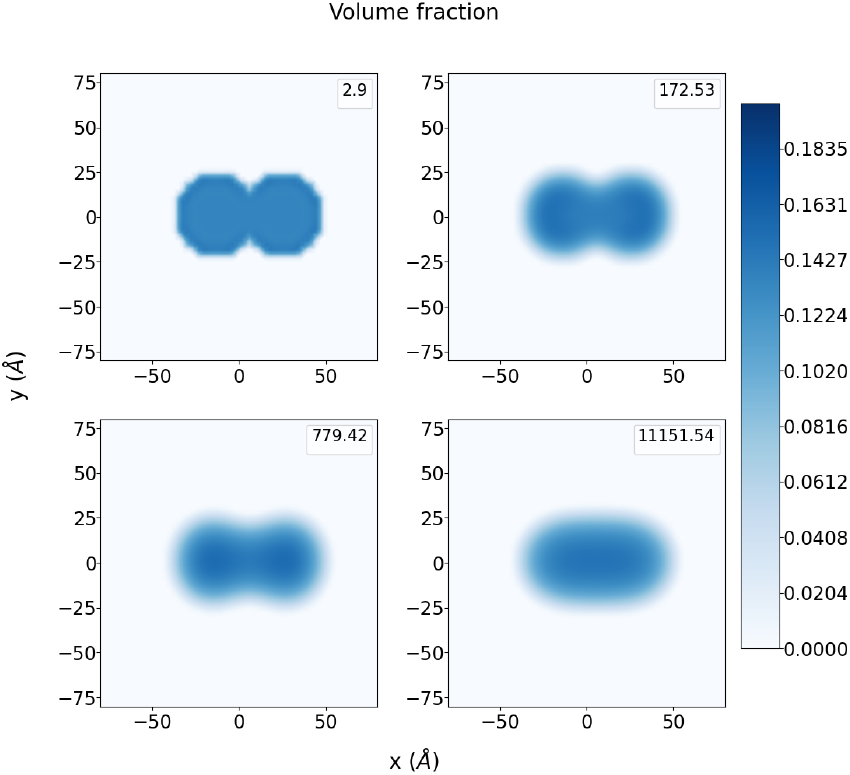
Time evolution of two coalescing droplets simulated in a 2D geometry with a simulation box of size 35nmx35nm (displayed snapshots from the simulation focus only on a portion of the full simulation box for visual clarity). The droplets are initialized with a volume fraction of φ = 0.127 and a dilute density volume fraction of φ = 3.8 · 10^−4^. The influence parameter is *κ* = 20. The color bar indicates the monomer volume fraction, and the timestamps in the top right of each panel are in units of *τ*_*s*_.

**FIG. 2.**
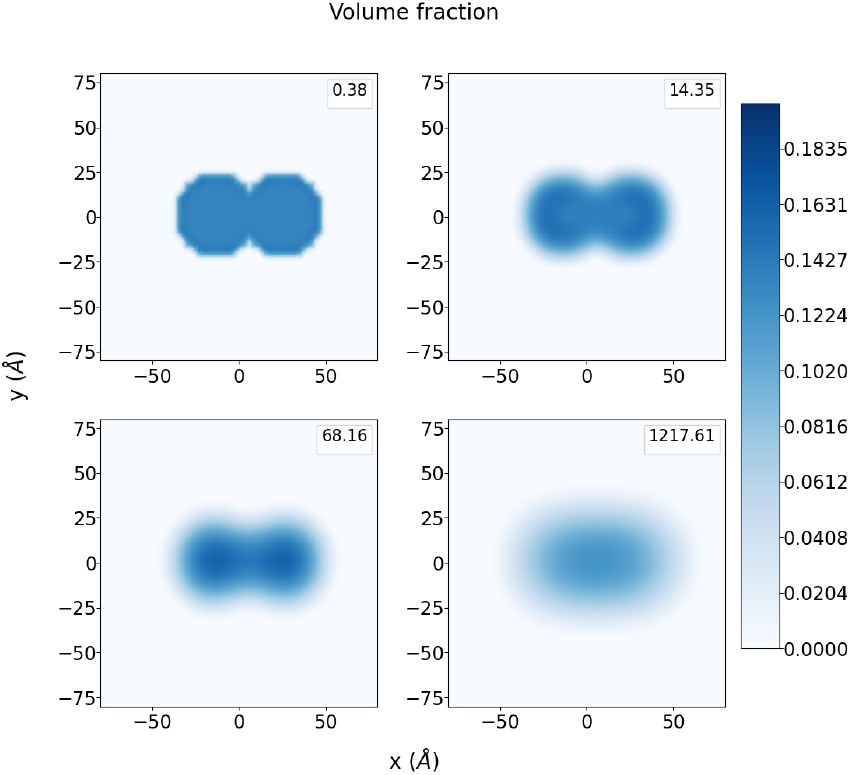
Time evolution of two coalescing droplets with higher influence parameter *κ* = 200. All other conditions are identical to Fig. 1. Note the significantly faster coalescence time compared to Fig. 1.

In both cases, coalescence proceeds via two stages: (i) densification of the individual droplets toward equilibrium, followed by (ii) physical fusion into a single larger droplet.

Stage (i) occurs rapidly and is driven by local chemical potential gradients. Stage (ii), which involves reorganization of the interface, proceeds more slowly and is governed primarily by surface tension forces. We argue that this observation is also consistent with all energy terms contributing to the dynamics of the first stage, whereas surface tension becomes the leading driving force in the second stage as the other energy terms approach equilibrium.

To quantify the influence of *κ*, we calculated the time to coalescence across a range of surface tension values. As shown in Fig. 3, coalescence time is inversely proportional to *κ* (*T*_*coalescence*_ ∝ 1*/κ*. This scaling is consistent with analytical models of droplet coalescence under Laplace pressure gradients^49–51^. Surface tension is the energetic cost of maintaining an interface between dense and dilute liquid phases; thus, larger *κ*, providing a stronger attraction for coalescing droplets, yields shorter coalescence times. Therefore, as the surface tension parameter *κ* increases, the time required for coalescence decreases.

**FIG. 3.**
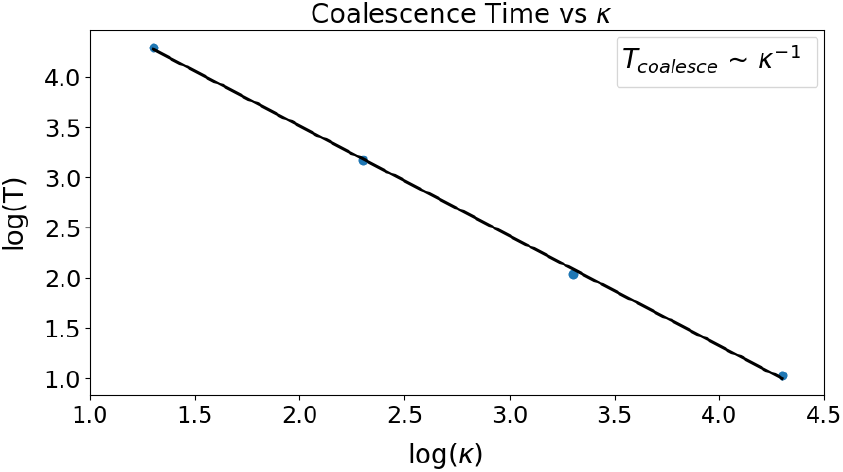
Coalescence time (*T*_*coalesce*_) as a function of the influence parameter *κ* for two droplets placed next to each other in 2D system. The log-log plot indicates inverse power-law relationship,*T*_coalesce_ ∼*κ*^−1^, indicating that stronger surface tensions leads to faster coalescence.

We also observe that the slope is nearly -1, which we attributed to the simulation’s 2D geometry, in which coalescence is driven by the merging of two circular disk droplets along the contact arcs. This finding validates the interpretation that surface tension dominates coalescence dynamics in the absence of hydrodynamics. We note that this scaling may be modified in 3D, which will be examined in future work.

Next, we investigated how an isolated underdense droplet relaxes toward equilibrium. We initialized single circular droplets with a uniform, step-like density embedded in a dilute phase at its equilibrium density. This setup mimics a single-phase domain that has not yet equilibrated its internal density profile or interface.

Fig. 4 shows the one-dimensional cross-sectional density profiles over time for two values of the surface tension parameter, *κ* = 20 (left) and *κ* = 200 (right), in a 115 nm × 115 nm simulation box. Over time, the sharp interface between the dense and dilute regions relaxes into a smoother profile as the system minimizes the total free energy. The interior of the droplet also becomes more homogeneous, eliminating the artificial flat-top density imposed initially. For lower *κ*, the relaxation is slower and the final profile retains a slightly flattened core. For higher *κ*, corresponding to larger interfacial energy penalties, the transition to a smooth profile is more rapid and results in a rounder, more compact droplet.

**FIG. 4.**
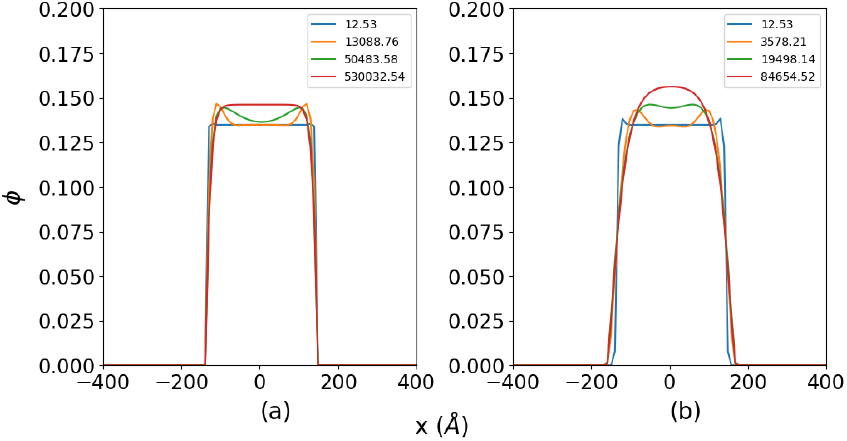
Single droplet (with initial uniform density of φ = 0.127) relaxation in a 115nm x115nm simulation box with *κ* = 20 (left) and *κ* = 200 (right). The surrounding dilute phase has a density of φ = 3.8 · 10^−4^.

To assess the sensitivity of our results to the model parameters, we repeated the coalescence simulations using an alternative set of Flory-Huggins parameters from Brady et al.^52^, corresponding to a different monomer to solvent volume ratio (*r*_*m*_ = 3.72). We also recalculated the influence parameter under these conditions, using Eq. 7, yielding *κ* = 161. The resulting dynamics are qualitatively similar (Fig. 5), although the timescales shift modestly due to parameter differences.

**FIG. 5.**
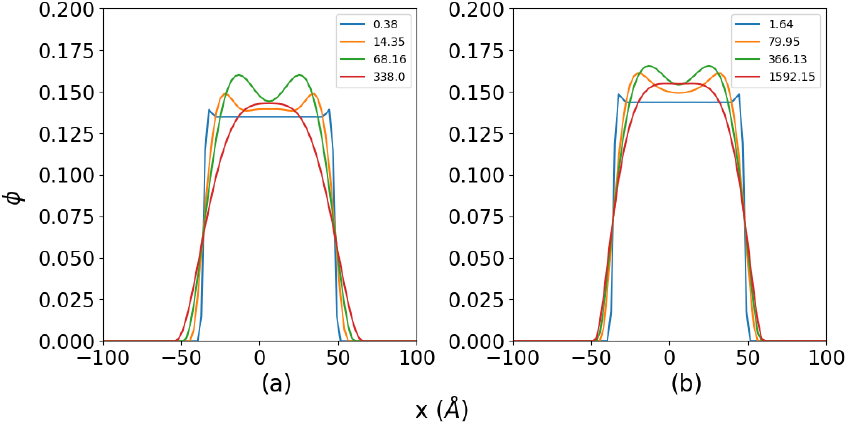
Comparison of droplet relaxation dynamics using different model parameters in a 29nm x 29 nm simulation box. DDFT simulation with *κ* = 200 and augmented Flory-Huggins parameters *ε*_*h*_ = 0.15, *ε*_*s*_ = −0.3 (left) and (b) Simulation with *κ* = 161, Flory-Huggins parameters *ε*_*h*_ = 0.2, *ε*_*s*_ = −0.3 . The droplet are initialized as a circular disk profile with φ = 0.127 and surrounded by more dilute phase with φ = 3.8 · 10^−4^

These simulations illustrate the capability of our DDFT framework to track both interfacial smoothing and internal density equilibration on mesoscale timescales. We note that the parameter space is vast. Our model includes several parameters, including those related to interfacial energy and molecular-scale interactions; comprehensive parameter sensitivity analyses are beyond the scope of this work. Our primary goal in this work is to demonstrate how the inclusion of a surface tension term within a physics-based DDFT framework enables the simulation of LLPS dynamics in dilute systems. Future efforts will more systematically explore parameter dependence, including both fitted and theoretically derived quantities, to further quantify the robustness and predictive range of the model.

We finally examined how coalescence and relaxation time scale with droplet diameter. Figure 6 shows a log-log plot of droplet relaxation and coalescence time *T* versus droplet diameter *D*, yielding slopes close to 3. The coalescence and relaxation timescale scaling as *T* ∝ *D*^3^ is consistent with surface-tension-limited dynamics. While DDFT does not incorporate hydrodynamics or viscous dissipation, this scaling mirrors that of hydrodynamic coalescence in inviscid surroundings, suggesting that droplet growth in our framework is governed by diffusive transport against surface tension gradients. The dilute surrounding phase, being several orders of magnitude less dense, effectively mimics an inviscid environment.

**FIG. 6.**
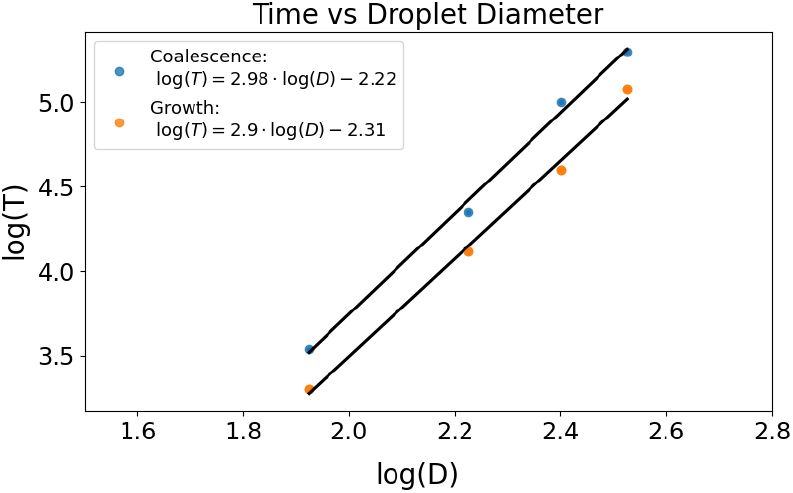
Scaling relationships for droplet relaxation time and coalescence time as a function of initial droplet diameter (D). The log-log plots show that both processes follow a power law with a slope close to 3. Extrapolating the coalescence data predicts that two droplets with a biologically relevant diameter of 2µm would coalesce in approximately 20 seconds for *κ* = 20 and 2 seconds for *κ* = 200. ( It is important to note that this prediction is based on a model that neglects hydrodynamic interactions, which may become more significant at this length scale and could alter the coalescence timescale.)

## IV. CONCLUSIONS

In this work, we developed and applied a dynamical density functional theory (DDFT) framework to model the nonequilibrium phase dynamics of a representative IDP undergoing LLPS. Our approach incorporates a free energy functional that combines Flory–Huggins–like interaction parameters, RPA-based electrostatic interactions, and a surface tension term to capture interfacial effects. While the present implementation uses FH parameters fitted to experimental phase diagrams, the framework is inherently designed to integrate descriptors derived directly from atomistic or coarse-grained simulations, enabling a predictive, simulation-based link between molecular interactions and continuum-scale dynamics.

Using this model, we demonstrated that DDFT can resolve the long-timescale evolution of dense and dilute phases, including droplet relaxation and coalescence, which would not be possible using mesoscale particle-based simulations. This establishes DDFT as a scalable continuum approach for studying non-equilibrium phase separation and interfacial phenomena relevant to biomolecular condensates.

Looking ahead, several important extensions remain. Applying the framework to a broader set of IDPs with varying sequence features will help clarify how molecular properties influence macroscopic dynamical phase behavior. Moreover, adapting the model to multicomponent systems, such as protein–RNA mixtures, will provide a powerful tool for probing the interplay between multiple biomolecular species in condensate formation and regulation.

Together, these directions will move us toward a unified, molecularly informed continuum framework capable of predicting the equilibrium and non-equilibrium behavior of complex biomolecular condensates across scales.

## ACKNOWLEDGMENTS

This work was supported by funding from the Cancer Prevention and Research Institute of Texas (CPRIT) award RR220008, the Welch Foundation (Award E-2221 and Catalyst Center for Advanced Bioactive Materials Crystallization Award V-E-0001), and NSF CBET-2442006. The simulations presented in this work were performed using the computational resources provided by the Hewlett-Packard Enterprise Data Science Institute at the University of Houston. The authors also thank Hasan Zerze for valuable discussions.

## DATA AVAILABILITY STATEMENT

All the data that support the conclusions of this study are given in the main text or supporting information. The DDFT code is available from the corresponding author upon request.

## Appendix A Free Energy Functional

Here, we adapted a free energy functional that captures the essential contributions governing the phase behavior of charged polymer solutions, inspired by the work of Chan and co-workers^27^. The total free energy density per unit volume, *f*, consists of three primary terms:

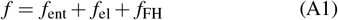

where *f*_ent_ describes the ideal mixing entropy, *f*_el_ accounts for electrostatic correlations via RPA, and *f*_FH_ represents shortrange monomer-solvent interactions using a Flory-Huggins-like form.

**1. Ideal Mixing Entropy**

We represented all molecular species on a lattice, with species *i* occupying a fraction φ_*i*_ of the total volume and composed of *N*_*i*_ monomers or segments. The mixing entropy per unit volume is:

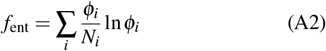

Here, φ_*i*_ = *ρ*_*i*_*a*^3^, where *ρ*_*i*_ is the number density of species *i*, and *a* is the lattice unit size. For polymers, *N*_*i*_ corresponds to chain length; for solvent or ions, *N*_*i*_ = 1.

**2. Electrostatic Correlations (RPA)**

To incorporate long-range electrostatic correlations beyond mean-field, we included a one-loop correction derived from RPA. The electrostatic free energy is given by:

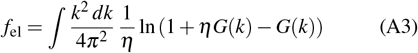

where *η* is the ratio of polymer segment volume to solvent molecular volume, and *G(k)* is the Fourier-transformed charge correlation function:

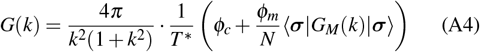

Here,

- 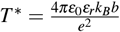 is the reduced (dimensionless) temperature.
- φ_*c*_ and φ_*m*_ are the volume fractions of counterions and monomers, respectively.
- ***σ*** is the vector of monomer charges (length *N*).
- *G*_*M*_*(k)*= is the intra-chain structure factor matrix, given by:

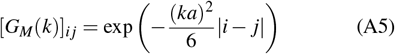

The term ⟨***σ*** |*G*_*M*_| ***σ***⟩ captures charge–charge correlations along the polymer chain, accounting for sequence specificity in electrostatic interactions.

**3. Short-Range Monomer–Solvent Interactions**

To model non-electrostatic short-range interactions, we included a Flory-Huggins-like enthalpic term:

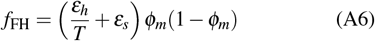

Here, *ε*_*h*_ and *ε*_*s*_ are effective parameters representing interaction enthalpy and entropy mismatch, respectively. In this work, we fit these parameters to match experimentally measured phase diagrams; however, they can also be extracted from simulation-derived coexistence curves.

The resulting free energy density (sum of all components in eqn. A1) allows us to capture essential physics of charged, phase-separating polymers—including entropy of mixing, sequence-specific electrostatics, and tunable short-range interactions—within a tractable continuum framework. This local free energy density is then converted to a global thermodynamic quantity, free energy functional, *F*^*′*^ as follows:

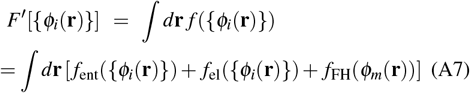

This integral over space yields a scalar-valued free energy functional that varies with the spatial profiles of all species. It serves as the thermodynamic driving potential in the DDFT evolution equation and allows for the computation of functional derivatives (chemical potentials) needed to describe the dynamics of each species.

## Appendix B Influence Parameter Derivation

In systems with dilute–dense coexistence, the two phases are separated by an interface of finite width. The excess free energy associated with this interface can be expressed as a surface tension, *γ*, multiplied by the interfacial area *S*:

In the dilute-dense phase coexistence, there is a surface tension, *γ*, between the two phases of contact area *S*, given by:

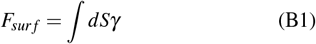

According to the density gradient model^37^, we can rewrite the surface energy in the form that depends on the gradient of density, such that the total energy can be written as:

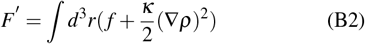

where *κ* is called the influence parameter. The surface tension can be derived by minimizing the Δ*F*^*′*^ with respect to *ρ*, where Δ*F*^*′*^ is given as:

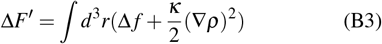

where Δ *f* = *f*-*ρf* ^⋆^(*ρ*), which is the difference between the Helmholtz free energy density of a homogeneous phase of density *ρ* and that of a two-phase equilibrium mixture ( *f*(^⋆^*ρ*)). In the case of a planar interface (1D) eqn B3 can be simplified as:

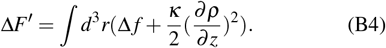

Minimizing eqn B4 with respect to density *ρ* results in density that satisfy this equation:

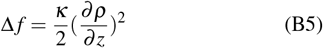

Finally, the surface tension can be written as:

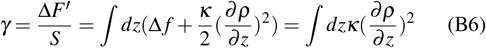

On the other hand, we know from eqn B5:

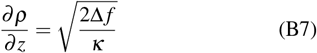

which turns equation B6 into :

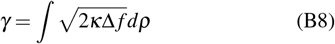

where the integral is done in *ρ* space from *ρ*_*dilute*_ inti *ρ*_*dense*_.

Furthermore, if we assume that *κ* is not function of *ρ*, we can write:

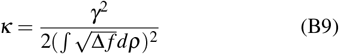

The term Δ *f* is given by:

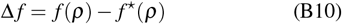

where:

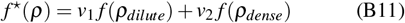

where *v*_1_ and *v*_2_ describe the volume fraction of dilute and dense phases.

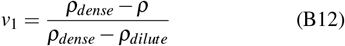

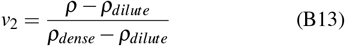

The term Δ *f* is always larger than 0 in [*ρ*_*dilute*_, *ρ*_*dense*_], because in this range, the liquid is more favorable in two-phase coexistence than in a uniform density.

## Notes

### Competing Interest Statement

The authors have declared no competing interest.

## References

1 S. F. Banani, H. O. Lee, A. A. Hyman, and M. K. Rosen, “Biomolecular condensates: organizers of cellular biochemistry,” Nature Reviews Molecular Cell Biology 18, 285–298 (2017).

2 Y. Shin and C. P. Brangwynne, “Liquid phase condensation in cell physiology and disease,” Science 357, eaaf4382 (2017).

3 S. Alberti, A. Gladfelter, and T. Mittag, “Considerations and challenges in studying liquid-liquid phase separation and biomolecular condensates,” Cell 176, 419–434 (2019).

4 S. Elbaum-Garfinkle, “Matter over mind: Liquid phase separation and neurodegeneration,” Journal of Biological Chemistry 294, 7160–7168 (2019).

5 B. Wang, L. Zhang, T. Dai, Z. Qin, H. Lu, L. Zhang, and F. Zhou, “Liquid– liquid phase separation in human health and diseases,” Signal transduction and targeted therapy 6, 290 (2021).

6 S. Boeynaems, S. Chong, J. Gsponer, L. Holt, D. Milovanovic, D. M. Mitrea, O. Mueller-Cajar, B. Portz, J. F. Reilly, C. D. Reinkemeier, et al., “Phase separation in biology and disease; current perspectives and open questions,” Journal of molecular biology 435, 167971 (2023).

7 A. Patel, L. Malinovska, S. Saha, J. Wang, S. Alberti, Y. Krishnan, and A. A. Hyman, “Atp as a biological hydrotrope,” Science 356, 753–756 (2017).

8 D. M. Mitrea, J. A. Cika, C. B. Stanley, A. Nourse, P. L. Onuchic, P. R. Banerjee, A. H. Phillips, C.-G. Park, A. A. Deniz, and R. W. Kriwacki, “Self-interaction of npm1 modulates multiple mechanisms of liquid–liquid phase separation,” Nature communications 9, 842 (2018).

9 L. Jawerth, E. Fischer-Friedrich, S. Saha, J. Wang, T. Franzmann, X. Zhang, J. Sachweh, M. Ruer, M. Ijavi, S. Saha, J. Mahamid, A. Hyman, and F. Julicher, “Protein condensates as aging maxwell fluids,” Science 370, 1317–1323 (2020).

10 J. A. Riback, L. Zhu, M. C. Ferrolino, M. Tolbert, D. M. Mitrea, D. W. Sanders, M.-T. Wei, R. W. Kriwacki, and C. P. Brangwynne, “Composition-dependent thermodynamics of intracellular phase separation,” Nature 581, 209–214 (2020).

11 T. S. Harmon, A. S. Holehouse, M. K. Rosen, and R. V. Pappu, “Intrinsically disordered linkers determine the interplay between phase separation and gelation in multivalent proteins,” elife 6, e30294 (2017).

12 G. L. Dignon, W. Zheng, Y. C. Kim, R. B. Best, and J. Mittal, “Sequence determinants of protein phase behavior from a coarse-grained model,” PLOS Computational Biology 14, e1005941 (2018).

13 J.-M. Choi, F. Dar, and R. V. Pappu, “Lassi: A lattice model for simulating phase transitions of multivalent proteins,” PLoS computational biology 15, e1007028 (2019).

14 J. A. Joseph, A. Reinhardt, A. Aguirre, P. Y. Chew, K. O. Russell, J. R. Espinosa, A. Garaizar, and R. Collepardo-Guevara, “Physics-driven coarsegrained model for biomolecular phase separation with near-quantitative accuracy,” Nature computational science 1, 732–743 (2021).

15 I. Yasuda, S. von Bulow, G. Tesei, E. Yamamoto, K. Yasuoka, and K. Lindorff-Larsen, “Coarse-grained model of disordered rna for simulations of biomolecular condensates,” Journal of chemical theory and computation 21, 2766–2779 (2025).

16 M. Paloni, R. Bailly, L. Ciandrini, and A. Barducci, “Unraveling molecular interactions in liquid-liquid phase separation of disordered proteins by atomistic simulations,” Journal of Physical Chemistry B 124, 9009–9016 (2020).

17 V. Ramachandran, W. Brown, C. Gayvert, and D. A. Potoyan, “Nucleoprotein phase-separation affinities revealed via atomistic simulations of short peptide and rna fragments,” The journal of physical chemistry letters 15, 10811–10817 (2024).

18 G. L. Dignon, W. Zheng, and J. Mittal, “Simulation methods for liquid– liquid phase separation of disordered proteins,” Current opinion in chemical engineering 23, 92–98 (2019).

19 P. J. Flory, “Thermodynamics of high polymer solutions,” The Journal of chemical physics 10, 51–61 (1942).

20 M. L. Huggins, “Some properties of solutions of long-chain compounds.” The Journal of Physical Chemistry 46, 151–158 (1942).

21 J. W. Cahn and J. E. Hilliard, “Free energy of a nonuniform system. i. interfacial free energy,” The Journal of chemical physics 28, 258–267 (1958).

22 J. D. Wurtz and C. F. Lee, “Chemical-reaction-controlled phase separated drops: formation, size selection, and coarsening,” Physical review letters 120, 078102 (2018).

23 J. E. Henninger, O. Oksuz, K. Shrinivas, I. I. Cissé, A. K. Chakraborty, and R. A. Young, “Rna-mediated feedback control of transcriptional condensates charge balance of electrostatic interactions can account for rna feedback regulation in brief during the early steps of transcription initiation, nascent rnas stimulate transcriptional condensate formation, whereas the burst of rnas produced during elongation stimulates condensate dissolution,” Cell 184, 207–225.e24 (2021).

24 P. A. Sharp, A. K. Chakraborty, J. E. Henninger, and R. A. Young, “Rna in formation and regulation of transcriptional condensates,” RNA 28, 52–57 (2022).

25 J. Berry, S. C. Weber, N. Vaidya, M. Haataja, and C. P. Brangwynne, “Rna transcription modulates phase transition-driven nuclear body assembly,” Proceedings of the National Academy of Sciences 112, E5237–E5245 (2015).

26 D. Zwicker, A. A. Hyman, and F. Jülicher, “Suppression of ostwald ripening in active emulsions,” Physical Review E 92, 012317 (2015).

27 Y.-H. Lin and H. S. Chan, “Phase separation and single-chain compactness of charged disordered proteins are strongly correlated,” Biophysical Journal 112, 2043–2046 (2017).

28 S. Das, A. N. Amin, Y.-H. Lin, and H. S. Chan, “Coarse-grained residue-based models of disordered protein condensates: utility and limitations of simple charge pattern parameters,” Physical Chemistry Chemical Physics 20, 28558–28574 (2018).

29 Y.-H. Lin, T. H. Kim, S. Das, T. Pal, J. Wessén, A. K. Rangadurai, L. E. Kay, J. D. Forman-Kay, and H. S. Chan, “Electrostatics of salt-dependent reentrant phase behaviors highlights diverse roles of atp in biomolecular condensates,” Elife 13, RP100284 (2025).

30 Y. H. Lin, J. D. Forman-Kay, and H. S. Chan, “Sequence-specific polyampholyte phase separation in membraneless organelles,” Physical Review Letters 117, 178101 (2016).

31 M. V. Nguyen, K. Dolph, K. T. Delaney, K. Shen, N. Sherck, S. Köhler, R. Gupta, M. B. Francis, M. S. Shell, and G. H. Fredrickson, “Molecularly informed field theory for estimating critical micelle concentrations of intrinsically disordered protein surfactants,” Journal of Chemical Physics 159 (2023), 10.1063/5.0178910/2931518.

32 G. H. Fredrickson and K. T. Delaney, “Direct free energy evaluation of classical and quantum many-body systems via field-theoretic simulation,” Proceedings of the National Academy of Sciences 119, e2201804119 (2022).

33 Y. O. Popov, J. Lee, and G. H. Fredrickson, “Field-theoretic simulations of polyelectrolyte complexation,” (2007).

34 J. Wessén, S. Das, T. Pal, and H. S. Chan, “Analytical formulation and field-theoretic simulation of sequence-specific phase separation of proteinlike heteropolymers with short-and long-spatial-range interactions,” The Journal of Physical Chemistry B 126, 9222–9245 (2022).

35 S. Najafi, J. McCarty, K. T. Delaney, G. H. Fredrickson, and J.-E. Shea, “Field-theoretic simulation method to study the liquid–liquid phase separation of polymers,” in Phase-Separated Biomolecular Condensates: Methods and Protocols (Springer, 2022) pp. 37–49.

36 M. Te Vrugt and R. Wittkowski, “Perspective: New directions in dynamical density functional theory,” Journal of Physics: Condensed Matter 35, 041501 (2022).

37 G. T. Dee and B. B. Sauer, “The surface tension of polymer liquids,” Advances in Physics 47, 161–205 (1998).

38 P. Taylor, “Ostwald ripening in emulsions,” Advances in Colloid and Interface Science 75, 107–163 (1998).

39 H. Tanaka, “Viscoelastic phase separation,” Journal of Physics: Condensed Matter 12, R207 (2000).

40 A. J. Bray, “Theory of phase-ordering kinetics,” Advances in Physics 51, 481–587 (2002).

41 O. R. Quayle, “The parachors of organic compounds. an interpretation and catalogue,” Chemical Reviews 53, 439–589 (1953).

42 S. C. Ayirala and D. N. Rao, “Application of the parachor model to the prediction of miscibility in multi-component hydrocarbon systems,” Journal of Physics: Condensed Matter 16, S2177 (2004).

43 J. E. Guyer, D. Wheeler, and J. A. Warren, “Fipy: Partial differential equations with python,” Computing in Science Engineering 11, 6–15 (2009).

44 M. Sala, W. Spotz, and M. Heroux, “PyTrilinos: High-performance distributed-memory solvers for Python,” ACM Transactions on Mathematical Software (TOMS) 34 (2008).

45 P. Yue, C. Zhou, and J. J. Feng, “Sharp-interface limit of the cahn–hilliard model for moving contact lines,” Journal of Fluid Mechanics 645, 279–294 (2010).

46 P. Yue, J. J. Feng, C. Liu, and J. Shen, “A diffuse-interface method for simulating two-phase flows of complex fluids,” Journal of Fluid Mechanics 515, 293–317 (2004).

47 P. Seppecher, “Moving contact lines in the cahn-hilliard theory,” International journal of engineering science 34, 977–992 (1996).

48 M. Rubinstein and R. H. Colby, Polymer Physics, 1st ed. (Oxford University Press, Oxford, 2003).

49 J. D. Paulsen, J. C. Burton, and S. R. Nagel, “Viscous to inertial crossover in liquid drop coalescence,” Phys. Rev. Lett. 106, 114501 (2011).

50 J. Eggers, J. R. Lister, and H. A. Stone, “Coalescence of liquid drops,” Journal of Fluid Mechanics 401, 293–310 (1999).

51 D. G. A. L. Aarts, H. N. W. Lekkerkerker, H. Guo, G. H. Wegdam, and D. Bonn, “Hydrodynamics of droplet coalescence,” Phys. Rev. Lett. 95, 164503 (2005).

52 J. P. Brady, P. J. Farber, A. Sekhar, Y. H. Lin, R. Huang, A. Bah, T. J. Nott, H. S. Chan, A. J. Baldwin, J. D. Forman-Kay, and L. E. Kay, “Structural and hydrodynamic properties of an intrinsically disordered region of a germ cell-specific protein on phase separation,” Proceedings of the National Academy of Sciences of the United States of America 114, E8194–E8203 (2017).

53 J. McCarty, K. T. Delaney, S. P. Danielsen, G. H. Fredrickson, and J.-E. Shea, “Complete phase diagram for liquid–liquid phase separation of intrinsically disordered proteins,” The journal of physical chemistry letters 10, 1644–1652 (2019).

54 Y. Lin, J. McCarty, J. N. Rauch, K. T. Delaney, K. S. Kosik, G. H. Fredrickson, J.-E. Shea, and S. Han, “Narrow equilibrium window for complex coacervation of tau and rna under cellular conditions,” Elife 8, e42571 (2019).

55 M. Paloni, R. Bailly, L. Ciandrini, and A. Barducci, “Unraveling molecular interactions in liquid–liquid phase separation of disordered proteins by atomistic simulations,” The Journal of Physical Chemistry B 124, 9009–9016 (2020).

56 K. L. Saar, D. Qian, L. L. Good, A. S. Morgunov, R. Collepardo-Guevara, R. B. Best, and T. P. Knowles, “Theoretical and data-driven approaches for biomolecular condensates,” Chemical Reviews 123, 8988–9009 (2023).

57 S. Boeynaems, S. Alberti, N. L. Fawzi, T. Mittag, M. Polymenidou, F. Rousseau, J. Schymkowitz, J. Shorter, B. Wolozin, L. Van Den Bosch, et al., “Protein phase separation: a new phase in cell biology,” Trends in cell biology 28, 420–435 (2018).

58 P. E. Wright and H. J. Dyson, “Intrinsically disordered proteins in cellular signalling and regulation,” Nature Reviews Molecular Cell Biology 2015 16:1 16, 18–29 (2014).

59 A. A. Hyman, C. A. Weber, and F. Jülicher, “Liquid-liquid phase separation in biology,” Annual review of cell and developmental biology 30, 39–58 (2014).

60 C. P. Brangwynne, T. J. Mitchison, and A. A. Hyman, “Active liquidlike behavior of nucleoli determines their size and shape in xenopus laevis oocytes,” Proc. Natl. Acad. Sci. 108, 4334–4339 (2011).

61 M. Kundu and M. P. Howard, “Dynamic density functional theory for drying colloidal suspensions: Comparison of hard-sphere free-energy functionals,” Journal of Chemical Physics 157 (2022), 10.1063/5.0118695/2842324.

62 K. Mazarakos, S. Qin, and H. X. Zhou, “Calculating binodals and interfacial tension of phase-separated condensates from molecular simulations with finite-size corrections,” Methods in Molecular Biology 2563, 1–35 (2023).

63 S. Qin and H. X. Zhou, “Fast method for computing chemical potentials and liquid-liquid phase equilibria of macromolecular solutions,” Journal of Physical Chemistry B 120, 8164–8174 (2016).

64 C. P. Brangwynne, C. R. Eckmann, D. S. Courson, A. Rybarska, C. Hoege, J. Gharakhani, F. Jülicher, and A. A. Hyman, “Germline p granules are liquid droplets that localize by controlled dissolution/condensation,” Science 324, 1729–1732 (2009).

65 M. T. Wei, Y. C. Chang, S. F. Shimobayashi, Y. Shin, A. R. Strom, and C. P. Brangwynne, “Nucleated transcriptional condensates amplify gene expression,” Nature Cell Biology 2020 22:10 22, 1187–1196 (2020).

66 S. Maharana, J. Wang, D. K. Papadopoulos, D. Richter, A. Pozniakovsky, I. Poser, M. Bickle, S. Rizk, J. Guillén-Boixet, T. M. Franzmann, M. Jahnel, L. Marrone, Y. T. Chang, J. Sterneckert, P. Tomancak, A. A. Hyman, and S. Alberti, “Rna buffers the phase separation behavior of prion-like rna-binding proteins,” Science (New York, N.Y.) 360, 918 (2018).

67 K. Gasior, M. G. Forest, A. S. Gladfelter, and J. M. Newby, “Modeling the mechanisms by which coexisting biomolecular rna–protein condensates form,” Bulletin of Mathematical Biology 82, 1–16 (2020).

68 N. Galvanetto, M. T. Ivanović, A. Chowdhury, A. Sottini, M. F. Nüesch, D. Nettels, R. B. Best, and B. Schuler, “Extreme dynamics in a biomolecular condensate,” Nature 2023 619:7971 619, 876–883 (2023).

69 A. Abyzov, M. Blackledge, and M. Zweckstetter, “Conformational dynamics of intrinsically disordered proteins regulate biomolecular condensate chemistry,” Chemical Reviews 122, 6719–6748 (2022).

70 A. Trifan, D. Gorgun, M. Salim, Z. Li, A. Brace, M. Zvyagin, H. Ma, A. Clyde, D. Clark, D. J. Hardy, T. Burnley, L. Huang, J. McCalpin, M. Emani, H. Yoo, J. Yin, A. Tsaris, V. Subbiah, T. Raza, J. Liu, N. Trebesch, G. Wells, V. Mysore, T. Gibbs, J. Phillips, S. C. Chennubhotla, I. Foster, R. Stevens, A. Anandkumar, V. Vishwanath, J. E. Stone, E. Tajkhorshid, S. A. Harris, and A. Ramanathan, “Intelligent resolution: Integrating cryo-em with ai-driven multi-resolution simulations to observe the severe acute respiratory syndrome coronavirus-2 replication-transcription machinery in action,” International Journal of High Performance Computing Applications 36, 603–623 (2022).

71 H. Lee, M. Turilli, S. Jha, D. Bhowmik, H. Ma, and A. Ramanathan, “Deepdrivemd: Deep-learning driven adaptive molecular simulations for protein folding,” Proceedings of DLS 2019: Deep Learning on Supercomputers - Held in conjunction with SC 2019: The International Conference for High Performance Computing, Networking, Storage and Analysis, 12–19 (2019).

72 H. I. Ingolfsson, C. Neale, T. S. Carpenter, R. Shrestha, C. A. Lopez, T. H. Tran, T. Oppelstrup, H. Bhatia, L. G. Stanton, X. Zhang, S. Sundram, F. D. Natale, A. Agarwal, G. Dharuman, S. I. K. Schumacher, T. Turbyville, G. Gulten, Q. N. Van, D. Goswami, F. Jean-Francois, C. Agamasu, D. Chen, J. J. Hettige, T. Travers, S. Sarkar, M. P. Surh, Y. Yang, A. Moody, S. Liu, B. C. van Essen, A. F. Voter, A. Ramanathan, N. W. Hengartner, D. K. Simanshu, A. G. Stephen, P. T. Bremer, S. Gnanakaran, J. N. Glosli, F. C. Lightstone, F. McCormick, D. V. Nissley, and F. H. Streitz, “Machine learning–driven multiscale modeling reveals lipid-dependent dynamics of ras signaling proteins,” Proceedings of the National Academy of Sciences of the United States of America 119, e2113297119 (2022).

73 H. Bhatia, T. S. Carpenter, H. I. Ingólfsson, G. Dharuman, P. Karande, S. Liu, T. Oppelstrup, C. Neale, F. C. Lightstone, B. V. Essen, J. N. Glosli, and P. T. Bremer, “Machine-learning-based dynamic-importance sampling for adaptive multiscale simulations,” Nature Machine Intelligence 2021 3:5 3, 401–409 (2021).

74 P. Tarazona and U. M. B. Marconi, “Beyond the dynamic density functional theory for steady currents: Application to driven colloidal particles in a channel,” Journal of Chemical Physics 128 (2008), 10.1063/1.2904881/984830.

75 M. te Vrugt, H. Löwen, and R. Wittkowski, “Classical dynamical density functional theory: from fundamentals to applications,” Advances in Physics 69, 121–247 (2020).

76 A. J. Archer, “Dynamical density functional theory for molecular and colloidal fluids: A microscopic approach to fluid mechanics,” Journal of Chemical Physics 130 (2009), 10.1063/1.3054633/902708.

77 B. Chun, T. Yoo, and H. W. Jung, “Temporal evolution of concentration and microstructure of colloidal films during vertical drying: a lattice boltzmann simulation study,” Soft Matter 16, 523–533 (2020).

78 R. Song, M. Lee, H. Moon, S. Lee, S. Shin, D. Kim, Y. Kim, B. Oh, and J. Lee, “Particle dynamics in drying colloidal solution using discrete particle method,” Flexible and Printed Electronics 6, 044007 (2021).

79 M. E. Cates, E. Tjhung, M. E. Cates, and E. Tjhung, “Theories of binary fluid mixtures: from phase-separation kinetics to active emulsions,” Journal of Fluid Mechanics 836, P1 (2018).

80 S. K. Das, J. Horbach, and K. Binder, “Kinetics of phase separation in thin films: Lattice versus continuum models for solid binary mixtures,” Physical Review E 79, 021602 (2009).

81 G. S. Grest, M. D. Lacasse, K. Kremer, and A. M. Gupta, “Efficient continuum model for simulating polymer blends and copolymers,” The Journal of Chemical Physics 105, 10583–10594 (1996).

82 G. H. Zerze, “Optimizing the martini 3 force field reveals the effects of the intricate balance between protein-water interaction strength and salt concentration on biomolecular condensate formation,” Journal of Chemical Theory and Computation 20, 1646–1655 (2024).

83 M. Beckinghausen and A. J. Spakowitz, “Interplay of polymer structure, solvent ordering, and charge fluctuations in polyelectrolyte solution thermodynamics,” Macromolecules 56, 136–152 (2023).

84 J. Holland, A. A. Castrejón-Pita, R. Tuinier, D. G. Aarts, and T. J. Nott, “Surface tension measurement and calculation of model biomolecular condensates,” Soft Matter 19, 8706–8716 (2023).

85 M. R. Mruzik, F. F. Abraham, and G. M. Pound, “Phase separation in fluid systems by spinodal decomposition. ii. a molecular dynamics computer simulation,” The Journal of Chemical Physics 69, 3462–3467 (1978).

86 G. H. Fredrickson and H. Orland, “Dynamics of polymers: A mean-field theory,” Journal of Chemical Physics 140 (2014), 10.1063/1.4865911/1005144.

87 M. Beckinghausen and A. J. Spakowitz, “Exploration of phase behavior in asymmetric semiflexible polyelectrolyte mixtures using polymer field theory,” Macromolecules 57, 2505–2519 (2024).

88 H. Kahl and S. Enders, “Calculation of surface properties of pure fluids using density gradient theory and saft-eos,” Fluid Phase Equilibria 172, 27–42 (2000).

89 W. Zheng, G. L. Dignon, N. Jovic, X. Xu, R. M. Regy, N. L. Fawzi, Y. C. Kim, R. B. Best, and J. Mittal, “Molecular details of protein condensates probed by microsecond long atomistic simulations,” Journal of Physical Chemistry B 124, 11671–11679 (2020).

90 J. A. Joseph, A. Reinhardt, A. Aguirre, P. Y. Chew, K. O. Russell, J. R. Espinosa, A. Garaizar, and R. Collepardo-Guevara, “Physics-driven coarse-grained model for biomolecular phase separation with near-quantitative accuracy,” Nature Computational Science 2021 1:11 1, 732–743 (2021).

91 H. Lin, Y. Y. Duan, and Q. Min, “Gradient theory modeling of surface tension for pure fluids and binary mixtures,” Fluid Phase Equilibria 254, 75–90 (2007).

92 D. Guo, Y. Xiong, B. Fu, Z. Sha, B. Li, and H. Wu, “Liquid-liquid phase separation in bacteria,” Microbiological Research 281, 127627 (2024).

93 H. Falahati and A. Haji-Akbari, “Thermodynamically driven assemblies and liquid–liquid phase separations in biology,” Soft Matter 15, 1135–1154 (2019).

94 S. A. Hollingsworth and R. O. Dror, “Molecular dynamics simulation for all,” Neuron 99, 1129 (2018).

95 J. D. Forman-Kay, J. A. Ditlev, M. L. Nosella, and H. O. Lee, “What are the distinguishing features and size requirements of biomolecular condensates and their implications for rna-containing condensates?” RNA 28, 36 (2022).

96 M. Gil-Garcia, A. I. Benítez-Mateos, M. Papp, F. Stoffel, C. Morelli, K. Normak, K. Makasewicz, L. Faltova, F. Paradisi, and P. Arosio, “Local environment in biomolecular condensates modulates enzymatic activity across length scales,” Nature Communications 2024 15:1 15, 1–11 (2024).

97 K. L. Saar, D. Qian, L. L. Good, A. S. Morgunov, R. Collepardo-Guevara, R. B. Best, and T. P. Knowles, “Theoretical and data-driven approaches for biomolecular condensates,” Chemical Reviews 123, 8988–9009 (2023).

98 H. H. Schede, P. Natarajan, A. K. Chakraborty, and K. Shrinivas, “A model for organization and regulation of nuclear condensates by gene activity,” Nature Communications 2023 14:1 14, 1–15 (2023).

99 M. te Vrugt, H. Löwen, and R. Wittkowski, “Classical dynamical density functional theory: from fundamentals to applications,” Advances in Physics 69, 121–247 (2020).

100 J. Lee, Y. O. Popov, and G. H. Fredrickson, “Complex coacervation: A field theoretic simulation study of polyelectrolyte complexation,” The Journal of chemical physics 128 (2008).

