## Supporting Information for "A Dynamical Density Functional Theory Framework for Non-Equilibrium Phase Dynamics in Biomolecular Condensates"

‡*Current address: Radiation Oncology Department, UT MD Andersen Cancer Center*

### Supporting Text

#### Sequence of Ddx4$^{n1}$

MGDEDWEAEINPHMSSYVPIFEKDRYSGENGDNFNRTPASSSEMDDGPSRRDHFMKSGFA
SGRNFGNRDAGECNKRDNTSTMGGFGVGKSFGNRGFSNSRFEDGDSSGFWRESSND-
CEDNPTRNRGFSKRGGYRDGNNSEASGPYRRGGRGSFRGCRGGFGLGSPNNDLDPDECMQ
RTGGLFGSRRPVLSGTGNGDTSQSRSGSGSERGGYKGLNEEVITGSGKNSWKSEAEGGES
